# Arginine Methylation Regulates SARS-CoV-2 Nucleocapsid Protein Function and Viral Replication

**DOI:** 10.1101/2021.03.24.436822

**Authors:** Ting Cai, Zhenbao Yu, Zhen Wang, Chen Liang, Stéphane Richard

## Abstract

Viral proteins are known to be methylated by host protein arginine methyltransferases (PRMTs) playing critical roles during viral infections. Herein, we show that PRMT1 methylates SARS-CoV-2 nucleocapsid (N) protein at residues R95 and R177 within RGG/RG sequences. Arginine methylation of N protein was confirmed by immunoblotting viral proteins extracted from SARS-CoV-2 virions isolated by cell culture. We demonstrate that arginine methylation of N protein is required for its RNA binding capacity, since treatment with a type I PRMT inhibitor (MS023) or substitution of R95K or R177K inhibited interaction with the 5’-UTR of the SARS-CoV-2 genomic RNA. We defined the N interactome in HEK293 cells with or without MS023 treatment and identified PRMT1 and many of its RGG/RG substrates including the known interactor, G3BP1, and other components of stress granules (SG). Methylation of N protein at R95 regulates another function namely its property to suppress the formation of SGs. MS023 treatment or R95K substitution blocked N-mediated suppression of SGs. Also, the co-expression of methylarginine reader TDRD3 quenched N-mediated suppression of SGs in a dose-dependent manner. Finally, pre-treatment of VeroE6 cells with MS023 significantly reduced SARS-CoV-2 replication. With type I PRMT inhibitors being in clinical trials for cancer treatment, inhibiting arginine methylation to target the later stages of the viral life cycle such as viral genome packaging and assembly of virions may be an additional therapeutic application of these drugs.

## Introduction

The COVID-19 pandemic is caused by severe acute respiratory syndrome coronavirus 2 (SARS-CoV-2), a virus which belongs to the family *coronaviridae* of genus *betacoronavirus* and has a positive sense strand RNA genome of ∼30 kb ^1^. It contains two large overlapping open reading frames (ORF1a and ORF1b) and encodes four structural proteins, namely spike (S), envelope (E), membrane (M) and nucleocapsid (N) proteins as well as nine accessory proteins ^1^. ORF1a and ORF1b are further processed to generate 16 non-structural proteins (Nsp1-16). Among the viral proteins, N protein is the most abundant in the virions and is expressed at the highest levels in infected cells ^2^. Thus, its abundance, essential roles during infection, and immunogenic nature, makes the SARS-CoV-2 N protein a valuable target for developing new strategies to combat the COVID-19 pandemic ^3–5^.

N protein regulates different steps of the coronavirus life cycle ^2^. The primary role of betacoronavirus N protein is the packaging of the viral genome into helical ribonucleoprotein (RNP) complexes ^6^. It is also involved in RNA synthesis with components of the replicase at early stages of infection ^7, 8^. Betacoronavirus N protein has two conserved and independently folded structural domains, called the N-terminal RNA-binding domain and C-terminal dimerization domain (NTD, CTD) ^4, 9, 10^, separated by flexible intrinsically disordered regions (IDRs) at the N-terminus, central serine/arginine-rich (SR) linker region, and C-terminal tail, respectively. The crystal structure of the SARS-CoV-2 NTD RNA binding domain depicts a U-shaped β-platform containing five short β-strands and an extended hairpin, forming a palm and finger-like structure with a highly positively charged cleft ^4, 11^.

After viral infection, host cells generate stress granules (SG) as an anti-viral response to inhibit protein synthesis and induce innate immune signaling ^12, 13^. SARS-CoV N protein plays an important role in host-virus interaction and localizes to cytoplasmic SGs ^14^. The SG nucleating factor G3BP1 ^15, 16^ and other SG components were identified in the SARS-CoV-2 N protein interactome ^5, 17^, suggesting SARS-CoV-2 like SARS-CoV regulates SGs mainly through N protein. The SARS-CoV-2 N protein is able to form condensates with RNA *in vitro* ^18–25^, in the cytoplasm of cells ^21, 25^ and partially co-localizes within arsenite-induced SGs ^26^. Several studies have shown that N protein sequesters G3BP1 and disassembles SGs ^22, 26–28^, likely as a means to suppress the host immune response to favor virus replication. Recent studies show that IDR1 and NTD regulate N protein condensates affecting nucleic acid annealing and potentially implicated in viral packaging and assembly ^21, 22, 25^. The SR linker region is phosphorylated by SRPK ^29^, GSK-3 ^30^ and Cdk1-GSK3 ^18^, influencing N protein condensates ^18, 22^. Besides phosphorylation, post-translational modifications that regulate N function are not known.

We identify that SARS-CoV-2 N protein contains five undefined and uncharacterized RGG/RG motifs. RGG/RG motifs are prevalent in RBPs and play key roles in mediating protein-protein and protein-RNA interactions ^31, 32^. The arginine residues located within the RGG/RG motifs are the preferred sites of methylation by protein arginine methyltransferases (PRMTs) ^33^. In mammals, there are nine PRMTs (PRMT1–9) that are classified into three types based on the methylmarks they produce: N^G^ -monomethylarginine (MMA), asymmetric N^G^, N^G^ - dimethylarginine (ADMA) and symmetric N^G^, Ń^G^ dimethylarginine (SDMA) ^33^. Methylarginines are bound by Tudor domains which are methylarginine readers ^34^. Arginine methylation regulates protein-protein interactions and protein-nucleic acid interactions to influence basic cellular processes, including transcription, RNA processing including pre-mRNA splicing, mRNA export, and mRNA translation, signaling transduction and liquid-liquid phase separation ^35, 36^. Unlike lysine demethylation, dedicated arginine demethylases have not been identified ^36^. Many specific small molecule inhibitors of PRMTs have been generated for cancer therapeutics ^37–42^ and a few have entered clinical trials (for review see ^36^).

Arginine methylation is known to methylate host and viral proteins necessary for viral replication. For example, the arginine methylation of HIV Tat protein decreases its transactivation function ^43^. The inhibition of PRMT5 prevents host hnRNPA1 RGG/RG motif methylation and inhibits HIV-1 and HTLV-1 IRES function ^44^. Moreover, PRMT5 methylates hepatitis B virus core (HBc) protein within its C-terminal arginine-rich domain to regulate its cellular localization ^45^. Arginine methylation of prototype foamy virus Gag in its glycine arginine rich box by PRMT1 regulates its nucleolar localization during replication ^46^.

In the present study, we report that PRMT1 methylates SARS-CoV-2 N protein within its RGG/RG motifs to regulate the RNA binding activity of the N protein towards its 5’-UTR genomic RNA. Moreover, arginine methylation modulates the role of N protein to inhibit SG formation. Our findings show for the first time that inhibition of type I PRMTs decreased SARS-CoV N methylation within virions and that arginine methylation is required for viral production.

## Results

### SARS-CoV-2 N protein is methylated by PRMT1

We noted that the SARS-CoV-2 N protein harbors two RGG (Figure 1A) and three RG motifs like SARS-CoV, but unlike MERS-CoV. As RGG/RG motifs are preferred sites of PRMT1, PRMT5 and PRMT6 ^31^, we tested whether they could be methylated. We first expressed and purified glutathione S-transferase (GST)-fusion proteins of the SARS-CoV-2 N protein fragments (GST-N 1-150, GST-N 150-262 and GST-N 263-419, Figure 1A) and performed *in vitro* arginine methylation assays. Both the N-terminal fragment (amino acid residues 1-150) and the central region (amino acid residues 150-262) were arginine methylated by PRMT1 (Figure 1B). By contrast, the N protein fragments were not methylated by PRMT5 or PRMT6 (Figure 1C and 1D). We then substituted arginines in the RGG/RG motifs to lysines to identify the methylated sites and maintain the charge. Mutation of arginine 68 (R68K) in the N-terminal fragment had no significant effect on arginine methylation, while mutation of arginine 95 (R95K) completely abolished PRMT1 methylation (Figure 1E), suggesting that R95 was the methylated residue in the GST-N 1-150 fragment. Similarly, mutation analysis identified R177 as the methylated residue in the central fragment (Figure 1F). Taken together, R95 and R177 within the RGG/RG motifs of the SARS-CoV-2 N were methylated *in vitro* by PRMT1.

**Figure 1.**
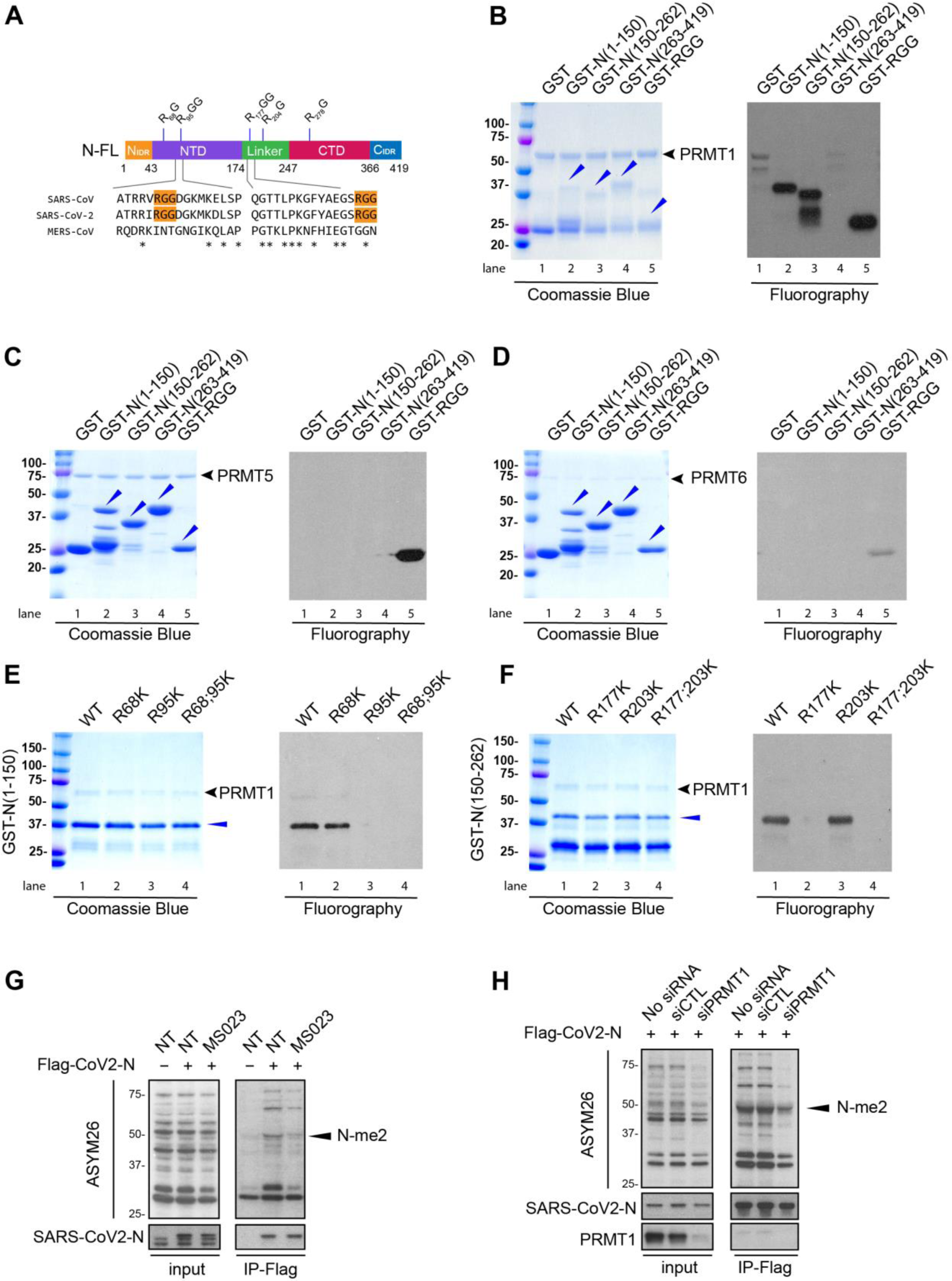
R95 and R177 within SARS-CoV-2 N RGG/RG motifs are methylated by PRMT1. (A) Schematic diagram of N protein with its N-terminal and C-terminal domains (NTD, CTD) and its N_IDR_ and C_IDR_ for N- and C-terminal intrinsic disordered regions and lastly the linker region between NTD and CTD known for its SR-rich sequences. Note R95GG and R177GG are conserved in SARS-CoV and SARS-CoV-2, but not MERS-CoV. (B, C, D) Recombinant GST-N protein fragments were subjected to *in vitro* methylation assays with recombinant B) GST-PRMT1, C) PRMT5/MEP50, and D) GST-PRMT6. Coomassie Blue staining and fluorography images are presented. GST alone and GST-RGG were used as negative and positive controls, respectively. Blue arrowheads indicate the migration of the GST-N protein fragments. The migration of GST-PRMT1, PRMT5 and GST-PRMT6 is shown on the right with a black arrow. The molecular mass markers are shown in kDa on the left. (E, F) GST-N protein fragments with arginine to lysine substitution were subjected to *in vitro* methylation assays. Coomassie Blue staining and fluorography images are presented. (G) HEK293 cells were transfected with control (-) or Flag-N (+) expression vectors for 24h and incubated with or without (NT) 1 μM MS023 for another 24h. The cell lysates were subjected to immunoprecipitation with anti-Flag-M2 beads and immunoblotting with anti-asymmetrical dimethylarginine antibody ASYM26 (upper panels) and anti-SARS-CoV-2 N protein antibody (lower panels). The band of the asymmetrically dimethylated N protein (N-me2) is marked by black arrowhead on the right. The molecular mass markers are shown in kDa on the left. (H) HEK293 cells were transfected with siRNA targeting firefly luciferase (siCTL) or siPRMT1 for 24h and subsequently transfected with Flag-N vector for another 24h. The cell lysates were subjected to immunoprecipitation with anti-Flag-M2 beads and then immunoblotting with anti-SARS-CoV-2 N protein and anti-ASYM26 antibodies. The migration of the methylated N protein is indicated.

We then determined whether the SARS-CoV-2 N protein was methylated in cells. HEK293 cells were transfected with a plasmid expressing Flag-epitope N protein (Flag-N). The cells were lysed and the N protein was immunoprecipitated with anti-Flag antibodies and its methylation detected by western blotting using the ASYM26 antibody which specifically recognizes asymmetrically dimethylated arginine (ADMA) residues within RGG/RG motifs. Importantly, the asymmetrical dimethylarginine methylation of the Flag-N protein (N-me2) was significantly reduced by treatment of the cells with the type I PRMT inhibitor MS023 (Figure 1G) and transfection with siPRMT1 (Figure 1H).

We next monitored patient data to identify modulation of PRMT1 expression during SARS-CoV-2 infection. Single cell RNA sequencing analysis of nasopharyngeal and bronchial samples from 19 clinically well-characterized SARS-CoV-2 patients and five healthy controls was performed ^47^. Importantly, analysis of their data showed that PRMT1 was significantly upregulated in infected patients (Figure S1). These data suggest PRMT1 may play a role during the SARS-CoV-2 life cycle.

### The SARS-CoV-2 N interactome defines a complex of RGG/RG proteins and PRMT1

We then performed mass spectrometry analysis to identify SARS-CoV-2 N-interacting proteins in the absence or presence of MS023. Flag-tagged SARS-CoV-2 N protein was expressed in HEK293 cells and a pull-down performed using anti-Flag affinity resin. Co-purified cellular proteins were subsequently analyzed by affinity-purification liquid chromatography and tandem mass spectrometry (AP-LC-MS/MS). We identified 119 cellular proteins interacting with SARS-CoV-2 N protein (peptide count > 2, fold change >2 between Flag-N and empty vector-transfected, 0.1% FDR, Figure 2A and B, Supplementary Dataset 1). Importantly, we identified several protein components of stress granule (SG) such as G3BP1 and G3BP2 (Ras-GTPase-activating protein SH3-domain-binding protein 1 and 2) ^48^, and CAPRIN1 (Figure 2A), in line with previous published AP-MS/MS studies ^5, 17^. Moreover, our mass spectrometry analysis identified SRSF protein kinase 1 (SRPK1) and GSK-3, known to phosphorylate N protein and regulate its localization to SGs ^18, 22, 29^. We also identified TRIM25 as a top hit in the N protein interactome. As a K63-linked ubiquitin ligase, TRIM25 mediates retinoic acid-inducible gene 1 (RIG-I) ubiquitination and activates TLR/RLR signaling pathway in response to RNA virus infection. It is known that SARS-CoV N protein interacts with TRIM25 and inhibits TRIM25/RIG-I association ^49^, suggesting that SARS-CoV-2 N protein may play a similar role in antagonizing the host immune response.

**Figure 2.**
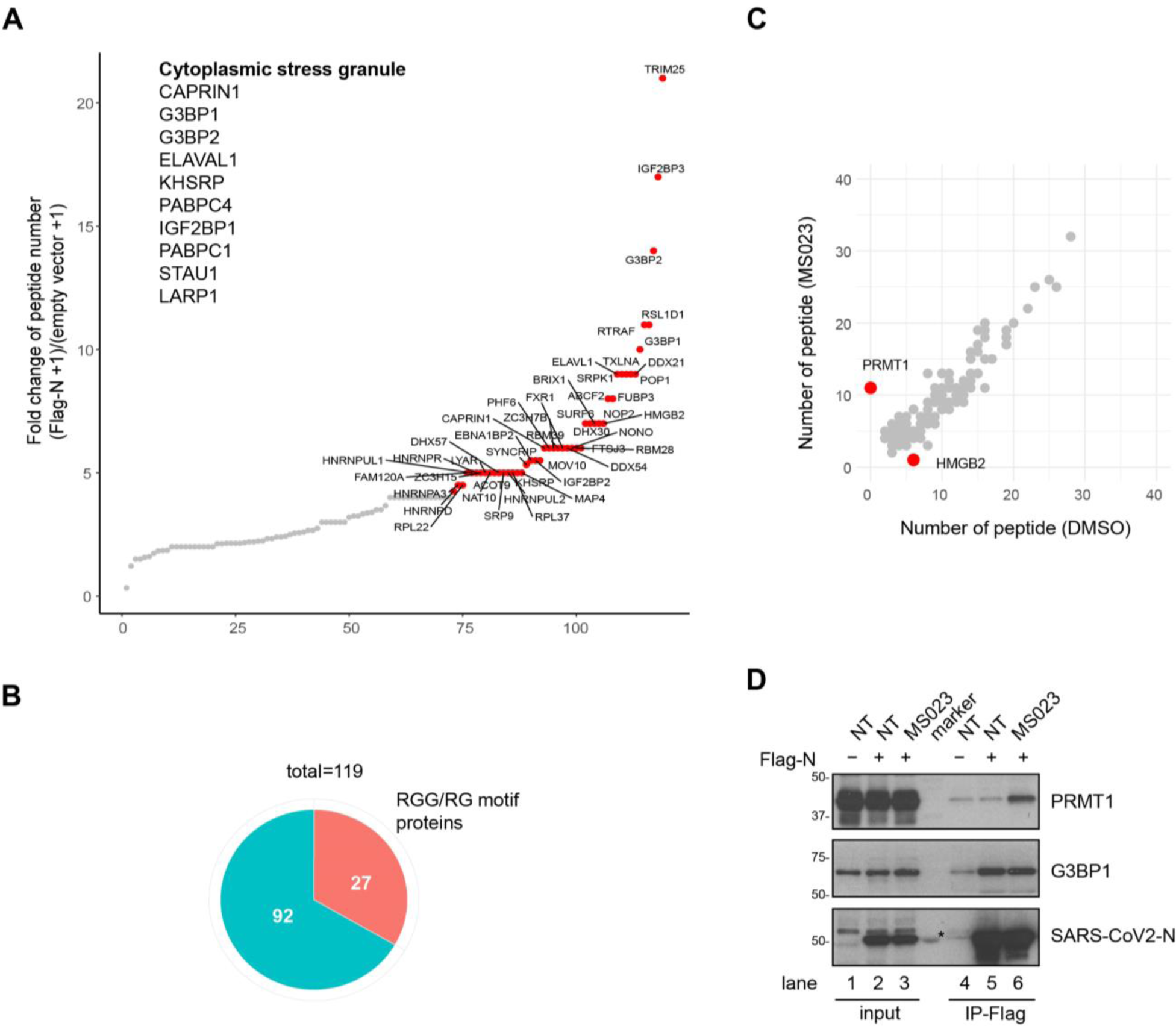
N protein interactome with and without MS023: association with many RGG/RG proteins and PRMT1. HEK293 cells were transfected with control or Flag-N and the next day Flag-N-transfected cells were subsequently treated with or without 1 μM MS023 for 24 h. Cell lysates were subjected to immunoprecipitation using anti-Flag-M2 beads. The bound proteins were identified by mass spectrometry (A-C). (A) Interactors were ranked by fold change of unique peptides detected from Flag-N-transfected cells and control plasmid-transfected cells (Flag-N + 1)/(empty vector + 1). Proteins with FC > 4 are highlighted in red. Immunoprecipitated proteins known to be localized in stress granule are listed. (B) Correlation analysis between MS023 and DMSO treated N protein interactome is shown. Proteins with a significant fold change (> 3 or < 3^-1^) after MS023 treatment are highlighted in red. (C) Pie chart represents the number of RGG/RG motif containing proteins among N protein interactors. (D) HEK293 cells were transfected with control (-) or Flag-N (+) and subsequentially treated with or without (NT) 1 μM MS023 for 24h. Cell lysates were immunoprecipitated with anti-Flag antibodies and the associated proteins separated by SDS-PAGE and immunoblotted with anti-PRMT1, anti-G3BP1 and anti-SARS-CoV-2 N antibodies. The asterisk denotes non-specific recognition of a molecular mass marker protein.

We then performed biological process (Gene Ontology) analysis using the identified interaction partners to assess major cellular pathways. The top 10 pathways enriched consisted of RNA metabolic processes (Figure S2). Interestingly, many SARS-CoV-2 N-interacting proteins (27 of 119, Figure 2B) contained multiple RGG/RG motifs including DEAD/DExH family of RNA helicases DDX21, DDX54, DHX30, DHX57, and heterogeneous nuclear ribonucleoproteins hnRNPA1, A3, D, DL, G (RBMX), R, U, UL1 and UL2 (Table 1). Many of these N-interacting proteins, such as G3BP1 ^50^, FAM98A ^51^, FXR1 ^52^, hnRNPA1 ^53^, hnRNPUL1 ^54^, SYNCRIP ^55^, ILF3 ^56^, and SERBP1^57^ are known PRMT1 substrates.

**Table 1.** RGG/RG motif containing proteins within SARS-CoV-2 N interactome.

| ID | Name | RNA binding |
| --- | --- | --- |
| Q14444 | CAPRIN1 | Yes |
| Q8TDD1 | DDX54 | Yes |
| Q6P158 | DHX57 | Yes |
| Q08211 | DHX9 | Yes |
| Q01844 | EWSR1 | Yes |
| Q9NZB2 | FAM120A | Yes |
| Q8NCA5 | FAM98A | Yes |
| P22087 | FBL | Yes |
| P51114 | FXR1 | Yes |
| Q13283 | G3BP1 | Yes |
| Q9UN86 | G3BP2 | Yes |
| O14979 | HNRNPDL | Yes |
| Q9BUJ2 | HNRNPUL1 | Yes |
| Q1KMD3 | HNRNPUL2 | Yes |
| Q14103 | HNRNPD | Yes |
| O60506 | SYNCRIP | Yes |
| O43390 | HNRNPR | Yes |
| Q00839 | HNRNPU | Yes |
| Q9NZI8 | IGF2BP1 | Yes |
| Q12905 | ILF2 | Yes |
| Q12906 | ILF3 | Yes |
| Q8NC51 | SERBP1 | Yes |
| Q99575 | POP1 | Yes |
| P38159 | RBMX | Yes |
| P09651 | HNRNPA1 | Yes |
| P51991 | HNRNPA3 | Yes |
| P46783 | RPS10 | Yes |

N protein-interactome changed significantly by two proteins with MS023 treatment. Interaction between N protein and HMGB2 was lost and interaction with PRMT1 was gained with MS023 treatment (Figure 2C). HMGB2 is a paralog of HMBG1 shown to play a critical role in SARS-CoV-2 replication ^58^. Interestingly, the AP-MS/MS analysis identified eleven PRMT1 peptides covering 30% of the protein sequence in the Flag-N immunoprecipitation from MS023-treated cells and none in non-treated cells (Figure 2C). These data are consistent with MS023 being non-competitive with type I PRMT substrates ^37^. To confirm these interactions, HEK293 cells were transfected with Flag-N and treated with DMSO control or MS023. Cell extracts were prepared and immunoprecipitations performed with anti-Flag antibodies followed by SDS-PAGE and immunoblotted with anti-PRMT1, -G3BP1 or -SARS-CoV2 N protein antibodies. Indeed, Flag-N immunoprecipitations showed increased PRMT1 association with MS023 treatment and G3BP1 was observed in immunoprecipitations from treated and non-treated cells (Figure 2D, compare lanes 5 and 6). These data confirm interactions between the N protein and G3BP1 and PRMT1.

### SARS-CoV-2 N prevents SG formation in an arginine methylation dependent manner

Stress granules are frequently observed upon infection with DNA or RNA viruses, serving an antiviral function ^12, 13^. Recent studies reveal that SARS-CoV-2 N protein is associated with SGs and regulates their dynamics ^22, 26–28^. To investigate whether arginine methylation regulates the property of SARS-CoV-2 N protein to suppress SG formation and dynamics, we monitored SG formation using anti-G3BP1 antibodies in the hepatoma Huh-7, a cell line frequently used in the study of SARS-CoV-2. Huh-7 cells transfected with an empty plasmid (pcDNA) or a plasmid expressing Flag-N protein were treated with a mild-dose of oxidative stress (0.5 μM sodium arsenite for 1h) or a harsh-dose (1 μM sodium arsenite for 2h). At 1 μM sodium arsenite, we observed Flag-N co-localizing with G3BP1 in SGs (open arrowheads) and some Flag-N expressing cells there was a reduction or absence of G3BP1 SGs (white arrowheads) (Figure 3A), as reported recently ^22, 26–28^. Interestingly, at the mild-doses of 0.5 μM sodium arsenite for 1 h, 25.89±2.56% of Flag-N protein transfected cells had G3BP1 SGs compared to 70.08±1.93% in the pcDNA transfected cells (Figure 3B), suggesting that SARS-CoV-2 N protein suppresses G3BP1 SG formation. As regulating SGs formation is an important function for viral replication and host cell immune response ^13^, we focused our study on how arginine methylation was implicated in N protein mediated SG suppression. Thus, all subsequent studies were performed with 0.5 μM sodium arsenite for 1 h to study N protein inhibition of G3BP1 SGs. Huh-7 cells transfected with Flag-N protein were treated with type I PRMT inhibitor MS023 or control DMSO before induction of SGs with sodium arsenite. Methyltransferase inhibition using MS023 significantly increased the presence of Flag-N expressing cells with G3BP1 SGs (45.48±4.79% versus 30.99±3.92%), while no significant change was observed in the non-transfected cells with over >70% of the cells with SGs (Figure 3C). PRMT1 inhibition in HeLa cells is known to increase the number of SGs per cell via RGG/RG motif methylation of G3BP1 ^50^ and UBAP2L^59^. To demonstrate the role of arginine methylation suppression of G3BP1 SGs was due to N protein methylation, *per se*, we transfected Huh-7 cells with wild type and R-K Flag-N proteins and monitored SGs. Cells with N protein with R95K or R95K/R177K substitution showed increased SG formation in comparison to those transfected with Flag-N or Flag-N R177K (Figure 4A, R95K 35.82±3.03%, R177K 26.74±2.52%, R95K/R177K 36.62±2.78% versus wild type N 25.30±2.62%). These findings show that the methylation of N protein at R95, but not R177, is required for the SARS-CoV-2 N to suppress G3BP1-positive SGs.

**Figure 3.**
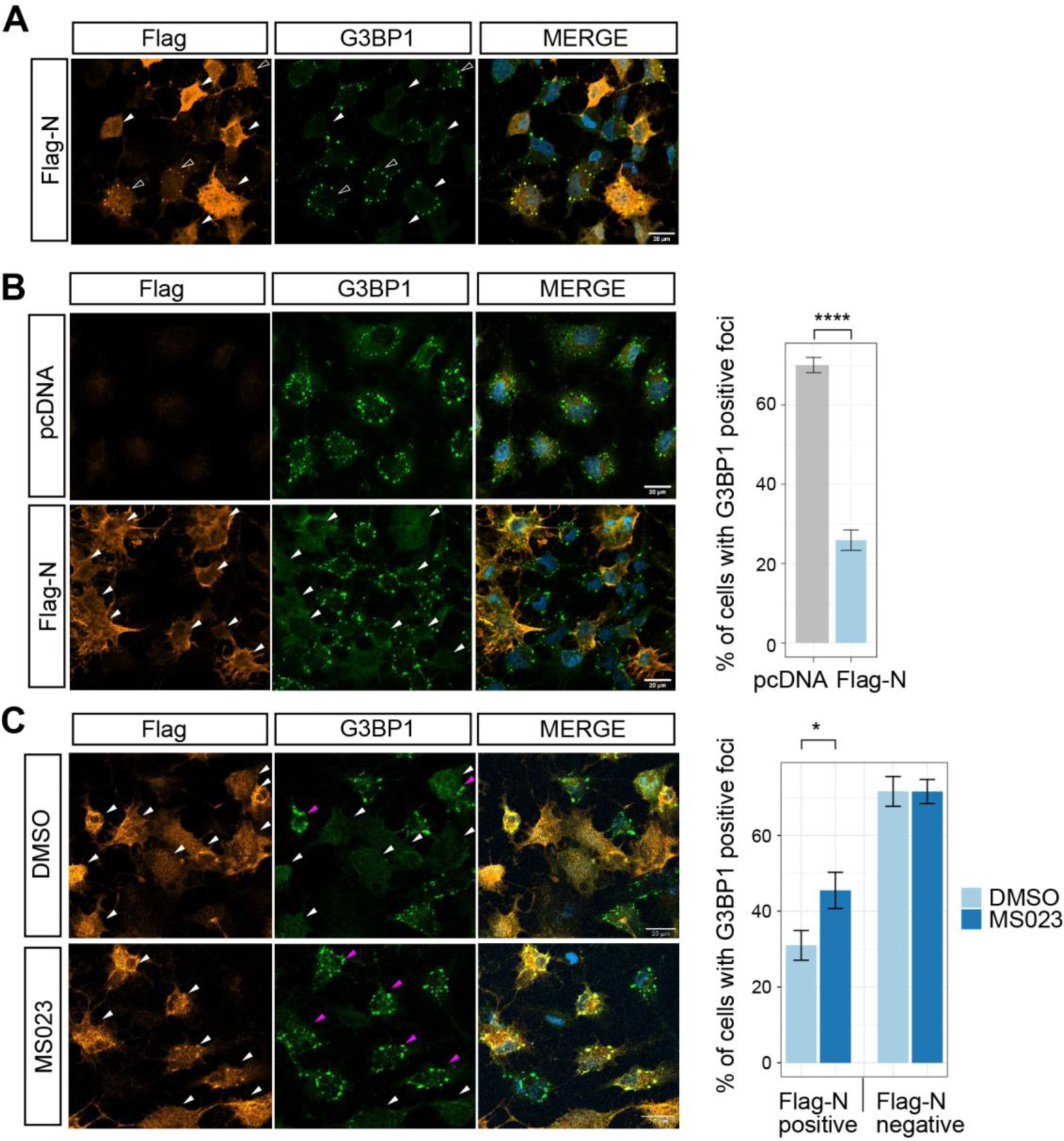
SARS-CoV-2 N protein regulates G3BP1 stress granule formation in an arginine methylation-dependent manner. (A) Huh-7 cells were transfected with control vector or Flag-N for 24h and subsequently incubated with 1 mM sodium arsenite for 2h. Cells were fixed with 4% PFA and co-immunostained with anti-Flag and anti-G3BP1 antibodies. A typical image is shown. Scale bar represents 20 μm. Arrowheads indicate FLAG-N-transfected cells, and the empty arrowheads indicate cells with FLAG-N and G3BP1 colocalization. (B) Huh-7 cells were transfected with control vector or Flag-N for 24h and subsequently incubated with 0.5 mM sodium arsenite for 1h. Cells were fixed with 4% PFA and co-immunostained with anti-Flag and anti-G3BP1 antibodies. A typical image is shown. Scale bar represents 20 μm. Arrowheads indicate FLAG-N-transfected cells, and transfected cells with SGs (> 5 G3BP1 foci) were highlighted in magenta. The percentage of cells harboring SGs (> 5 G3BP1 foci) were quantified and shown in the bar plot on the right. n =15 fields from 3 independent experiments are shown. Welch’s t test. \*\*\*\**p* < 0.0001. (C) Huh-7 cells were transfected with Flag-N overnight and treated with or without 5 μM MS023 for another 24h. Then the cells were incubated with 0.5 mM sodium arsenite for another hour and fixed with 4% PFA. Cells were co-immunostained with anti-Flag and anti-G3BP1 antibodies. A typical image is shown in the left panel. Scale bar, 20 μm. Arrowheads indicate Flag-N-transfected cells, and transfected cells with SGs (> 5 G3BP1 foci) were highlighted in magenta. Percentage of cells harboring SGs (> 5 G3BP1 foci) were quantified in the transfected cells (Flag-N positive) and non-transfected cells (Flag-N negative) respectively and shown in the bar plot on the right. n =20 fields from 2 independent experiments are shown. Welch’s t test. \**p* < 0.05.

**Figure 4.**
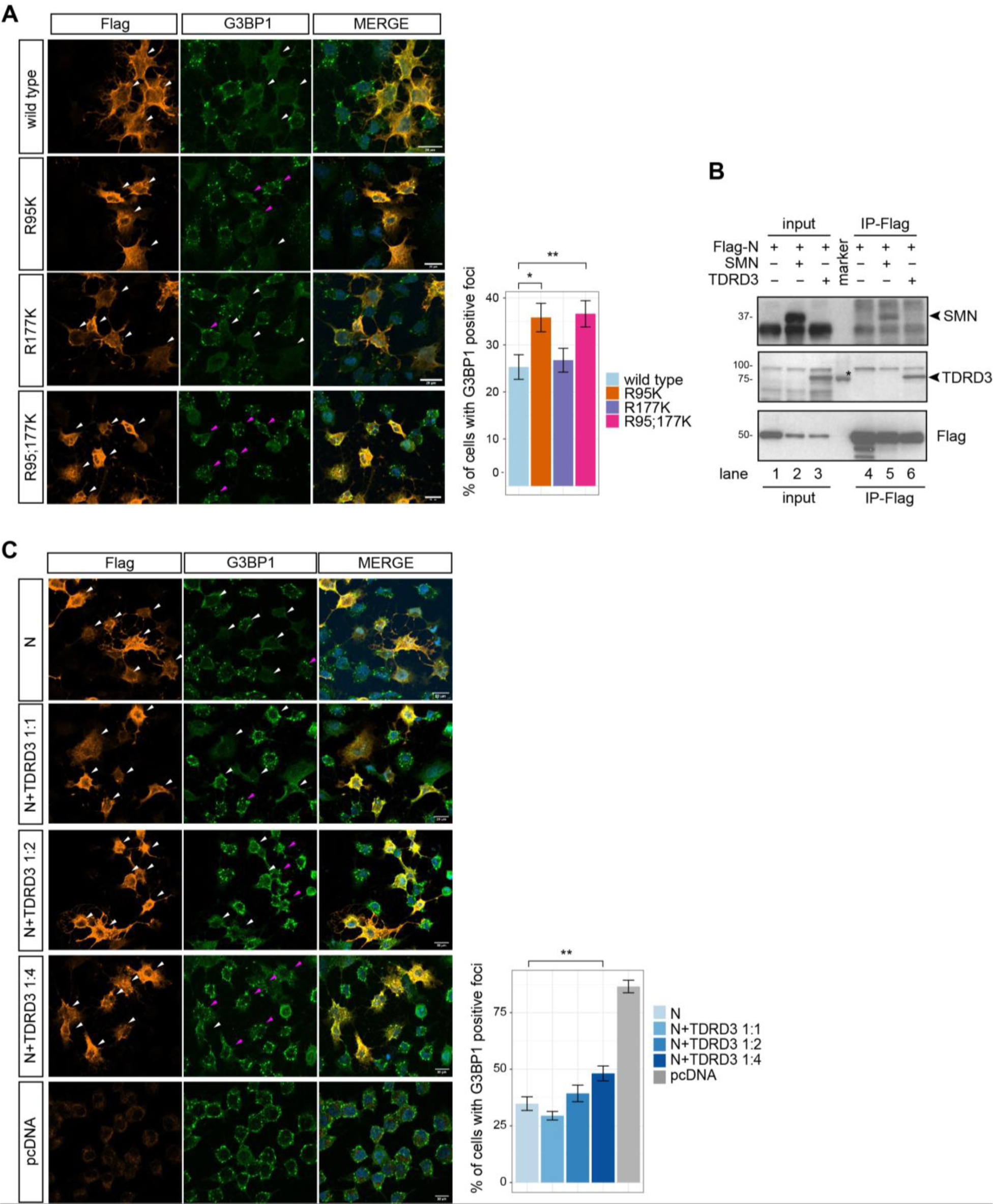
R95 is required for N protein regulating G3BP1 stress granule and methylarginine reader protein TDRD3 is involved in this process. (A) Huh-7 cells were transfected with Flag-N (wild type) and its mutants for 24h and subsequently incubated with 0.5 mM sodium arsenite for another hour. Cells were fixed with 4% PFA and co-immunostained with anti-Flag and anti-G3BP1 antibodies. A typical image is shown in the left panel. Scale bar, 20 μm. Arrowheads indicate transfected cells, and transfected cells with SGs (> 5 G3BP1 foci) were highlighted in magenta. Percentage of cells harboring SGs (> 5 G3BP1 foci) in the Flag-N-positive cell group were quantified and shown in the bar plot on the right. n =15 fields from 3 independent experiments are shown. Welch’s t test. \**p* < 0.05, \*\**p* < 0.01. (B) HEK293 cells were co-transfected with Flag-N and myc-SMN or myc-TDRD3. 24h later, cell lysates were immunoprecipitated with anti-Flag antibodies and the associated proteins separated by SDS-PAGE and immunoblotted with anti-SMN, anti-TDRD3 and anti-Flag antibodies. The asterisk denotes non-specific recognition of a molecular mass marker protein. (C) Huh-7 cells were co-transfected with Flag-N and myc-TDRD3 with indicated plasmid ratios. 24h later, cells were incubated with 0.5mM sodium arsenite for another hour. Cells were fixed with 4% PFA and co-immunostained with anti-Flag and G3BP1 antibodies. A typical image is shown on the left. Scale bar, 20 μm. Arrowheads indicate transfected cells, and transfected cells with SGs (> 5 G3BP1 foci) were highlighted in magenta. Percentage of the transfected cells (Flag-N positive) harboring SGs (> 5 G3BP1 foci) were quantified and shown in the bar plot on the right. n >15 fields from 2 independent experiments are shown. Welch’s t test. \*\**p* < 0.01.

We next examined whether the ectopic expression of a methylarginine reader could also interfere with N protein-mediated of G3BP1 SGs. The Tudor domain is a known reader of methylated arginine residues ^34^. We first examined whether Flag-N protein associated with ectopically expressed Tudor domain-containing proteins SMN and TDRD3 by coimmunoprecipitation assays. Indeed, a strong interaction between Flag-N protein and TDRD3 was observed, whereas the interaction with SMN was weaker, as visualized by immunoblotting (Figure 4B). Next, we tested whether TDRD3 influenced N protein-mediated SG regulation, as it is known that TDRD3 localizes to SGs ^60, 61^. We co-transfected increasing amounts of expression plasmid encoding TDRD3 together with Flag-N protein and visualized the presence of arsenite-induced G3BP1 SGs. We observed an increase in G3BP1 SGs with increase expression of TDRD3 with N protein (Figure 4C). These findings suggest that an increasing the methylreader TDRD3 expression could be a means to quench the effects of N protein on SG regulation.

### Arginine methylation of R95 and R177 is required for N protein binding to the SARS-CoV-2 5’-UTR RNA

N is an RNA binding protein that binds the 5’-UTR of its viral genomic RNA for viral ribonucleoprotein (vRNP) formation and packaging into virions ^62, 63^. The SARS-CoV-2 N protein R95 and R177 are located in the NTD domain and at NTD-SR linker boundary, respectively. Actually, R95 and R177 are within the RNA binding site of the NTD with R177 being predicted to be implicated in N protein RNA binding ^4, 11, 64^. Therefore, we reasoned that these arginines and their methylation were likely involved in the RNA binding activity of N protein. HEK293 cells were co-transfected with Flag-N and an expression vector transcribing ∼400 bp of the 5’-UTR RNA sequence of SARS-CoV-2 (p5’-UTR:CoV-2; Figure 5A). Initially, we performed RNA immunoprecipitation (RIP) to monitor N protein RNA binding activity. We observed a >5-fold enrichment with anti-Flag antibodies in the DMSO treated cells versus MS023 treated cells (Figure 5A). These data suggest that inhibition of type I PRMTs prevent the binding of Flag-N to the 5’-UTR of SARS-CoV-2 RNA. Next, we wished to confirm that this N protein/RNA interaction was direct by performing a photoactivatable ribonucleoside-enhanced crosslinking and immunoprecipitation (PAR-CLIP) assay ^65^. Cells transfected with Flag-N or Flag-N harboring R95K, R177K, R95K/R177K and p5’-UTR:CoV-2 were labeled with 4-thiouridine and UV cross-linked. The cells were lysed and immunoprecipitated with anti-Flag antibodies following a ‘Clipping’ step with RNase A. RNA was purified and analyzed by RT-qPCR with two sets of primers against the 5’-UTR of the SARS-CoV-2 RNA (position #1 and #2). Using this strategy, we showed that Flag-N directly binds to the 5’-UTR of SARS-CoV-2 RNA (Figure 5B). Importantly, both the single R95K and R177K or the double R95K/R177K substitution of N protein abolished RNA binding activity (Figure 5B). Immunoblotting was performed to confirm an equal expression of wild type N protein and the R-K proteins immunoprecipitated of the four replicates (Figure 5C). Taken together, these results suggest that arginine methylation of both R95 and R177 of the N protein is a requirement for association with its viral RNA *in cellulo*.

**Figure 5.**
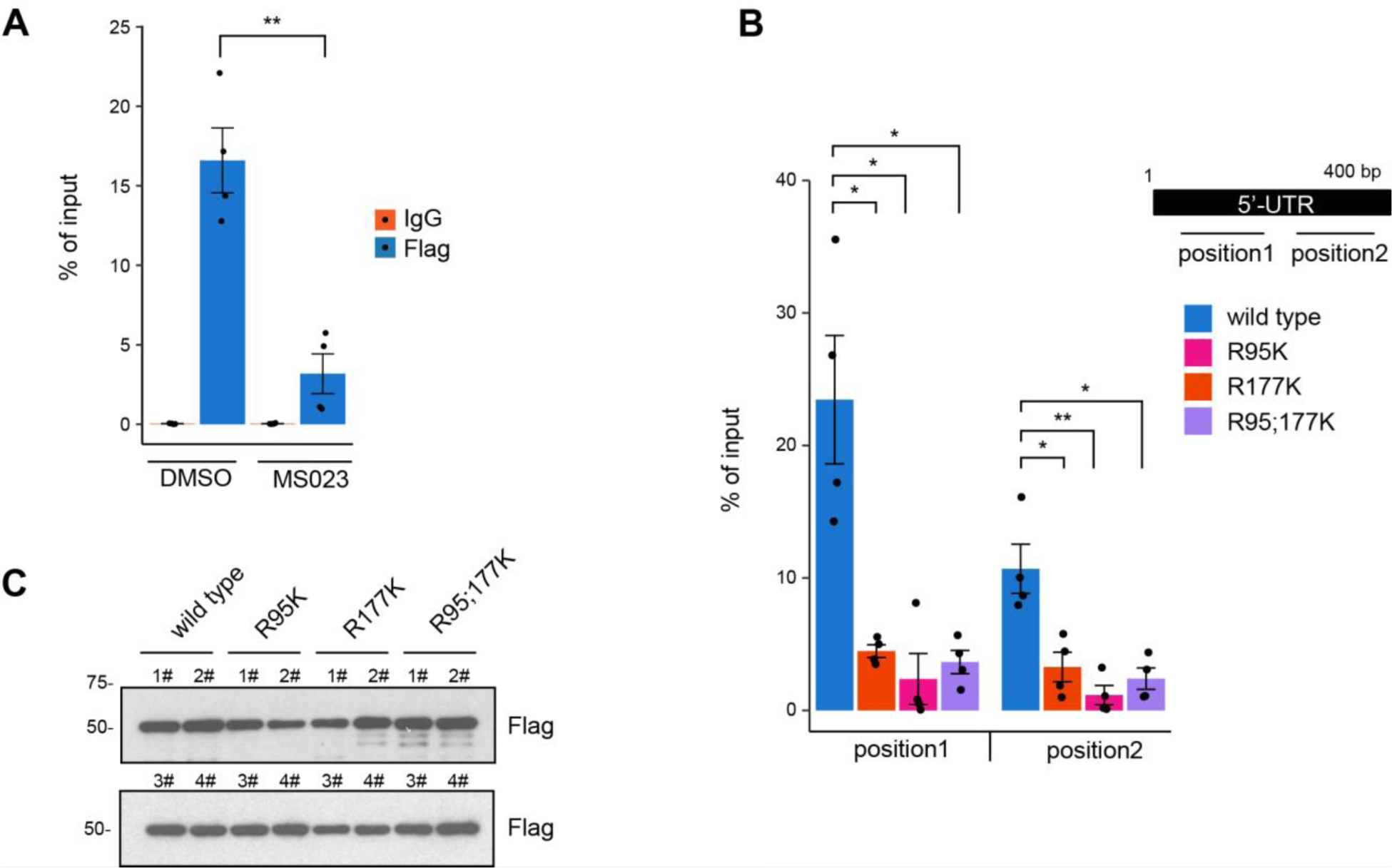
Arginine methylation of N R95 and R177 is a requirement for SARS-CoV-2 N binding to the 5’-UTR of its genomic RNA. (A) HEK293 cells were co-transfected with the plasmid expressing Flag-N protein and the plasmid expressing a 400 bp RNA fragment of the SARS-CoV-2 5’-UTR region (p5’-UTR:CoV-2, NC_045512: 1-400 bp). The cells were incubated with 5 μM MS023 or DMSO for 24 h. Cells were cross-linked with 1% formaldehyde and subjected to RIP using IgG or anti-Flag antibodies. The immunoprecipitated RNA was extracted and RT-qPCR was used to assess the bound RNA. Data is shown as percentage of input from 2 independent experiments. Welch’s t test. \*\**p* < 0.01. (B and C) HEK293 cells were co-transfected with plasmids expressing wild type and RK N proteins along with the plasmid expressing p5’-UTR:CoV-2 RNA. Then the cells were treated with 4-thiouridine for 16h and subjected to PAR-CLIP analysis (B). RT-qPCR with primers targeting consecutive regions, shown in the diagram, were used to assess the bound RNA. Data is shown as percentage of input from 2 independent experiments. Welch’s t test. \**p* < 0.05, \*\**p* < 0.01. Immunoblotting for expression of indicated proteins from 2 independent experiments is shown respectively in the top and bottom panels (C).

### Methylation of N protein is required for SARS-CoV-2 production

We then studied the effect of type I PRMT inhibitor MS0233 on SARS-CoV-2 replication. First, we performed MS023 toxicity assays with VeroE6 cells, a cell line highly susceptible to SARS-CoV-2 infection. We confirmed that cell proliferation was not affected at concentrations of MS023 up to 30 µM in complete medium (Figure 6A). Next in a certified SARS-CoV-2 BL3 laboratory, we then treated VeroE6 cells with 10 µM or 20 µM MS023, versus DMSO control, for 24 h and proceeded with SARS-CoV-2 infection at a low multiplicity of infection (MOI=0.1). The infected cells were kept in MS023 containing medium for another two days. Supernatant from each, infected well was collected and the virus inactivated with TRIzol to assay SARS-CoV-2 titer by Taqman real-time PCR assay. We observed a significant reduction of viral titer when the cells were incubated with 20 µM MS023 and an intermediate viral titer was observed with 10 µM MS023 (Figure 6B).

**Figure 6.**
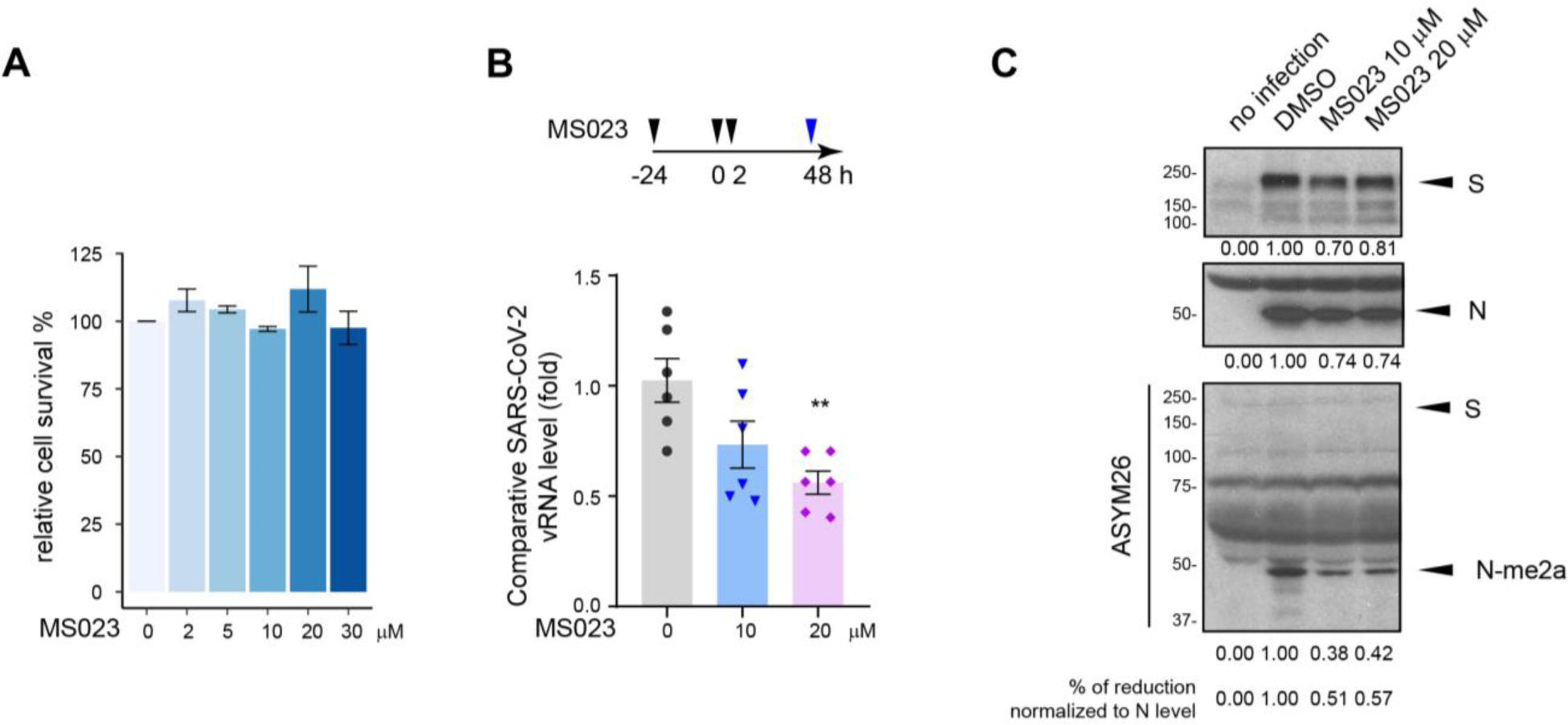
SARS-CoV-2 replication is impaired by type I PRMT inhibitor MS023. (A) VeroE6 cells were treated with the indicated dose of MS023 for 3 days as described in the “Materials and Methods” section and cell viability was determined by MTT assay. Data represents percentage of survival compared to control (DMSO alone) from 3 independent experiments. (B) VeroE6 cells were pretreated with MS023 or vehicle DMSO as control for 24 h. Then the cells were infected with SARS-CoV-2 at an MOI of 0.1. Two hours later, the medium was refreshed, and cells were incubated with the same concentration of MS023 or DMSO, respectively. Forty-eight hours later, the cell medium containing the released SARS-CoV-2 virions was collected and lysed in TRIzol immediately. Viral RNA was isolated and Taq-man probe base RT-qPCR was used to assess the viral load. Data represents the fold change against control samples from 2 independent experiments. Welch’s t test. \*\**p* < 0.01. (C) Viral proteins were extracted from the organic phase of samples in (B) and immunoblotted with anti-S, anti-N, and anti-ASYM26 antibodies. The density of the protein bands was calculated using ImageJ.

From the TRIzol organic phase, we extracted the viral proteins and separated them by SDS-PAGE followed by immunoblotting with anti-SARS-CoV-2 S, and N protein or anti-methylarginine ASYM26 antibodies. We observed a slight decrease (∼20-30%) in the total amounts of S and N proteins with MS023 treatment (Figure 6C), consistent with overall decrease in virions of (Figure 6B). Notably, the anti-ASYM26 antibodies revealed a ∼50% reduction in N protein arginine methylation, when normalized to N protein levels (Figure 6C, middle panel). In contrast, we could not detect methylation of S with ASYM26 antibody (Figure 6C), likely due to the lack of RGG/RG motifs in S protein sequence. These findings show that the SARS-CoV-2 production is reduced in the presence of type I PRMT inhibitors and that the SARS-CoV-2 N protein is arginine methylated within the virions.

## Discussion

In the present manuscript, we identify SARS-CoV-2 N protein to be arginine methylated within virions. We show that methylation of the N protein is mediated by PRMT1 at R95 within the NTD and R177 at the NTD/SR linker boundary. Both residues are within RGG/RG motifs conserved between SARS-CoV and SARS-CoV-2. Amino acid substitutions R95K or R177K inhibited N protein RNA binding *in cellulo* to SARS-CoV-2 5’-UTR genomic RNA using a CLIP assay in HEK293 cells. The ectopic expression of N protein in Huh-7 cells was localized to cytoplasmic granules and inhibition of G3BP1 stress granule (SG) formation was observed. Notably, arginine methylation of N protein at R95 by PRMT1 was necessary for this function. The N protein interactome was defined to contain many known PRMT1 substrates with RGG/RG motifs. Treatment with the type I PRMT inhibitor MS023 did not influence the overall N interactome, but PRMT1 was identified as a new interactor, consistent with a substrate-enzyme interaction in the presence of the non-competitive inhibitor MS023 ^37^. Importantly, inhibition of arginine methylation with MS023 significantly reduced SARS-CoV-2 replication in VeroE6 cells. Our findings define arginine methylation as new mode of interfering with N protein regulation of SGs and define PRMT1 as a requirement for the SARS-CoV-2 life cycle. As type I PRMT inhibitors are in clinical trials for cancer treatment ^41^, these compounds may also be useful to target SARS-CoV-2 replication.

Host proteins and enzymes are needed for the replication of SARS-CoV-2 and many of these were identified using CRISPR screens ^58, 66–69^. Factors required for early viral entry and fusion, and components of the endosome and cholesterol pathways were identified ^66–68^. Few components, however, were identified that target the later phases of the viral life cycle, such as vRNP formation, phase separation for genome packaging, encapsidation and assembly of virions. A recent study identified ∼50 host RBPs binding SARS-CoV-2 genomic RNA and the knockdown of some of these RBPs inhibited viral replication ^70^. As SARS-CoV-2 N protein and cellular RBPs are substrates of PRMT1 ^31^, our findings suggest that type I PRMT inhibitors or increasing methylarginine readers may regulate N function in vRNP packaging and assembly into virions. Notably, the increased expression of PRMT1 in nasopharyngeal and bronchial samples of SARS-CoV-2 infected patients ^47^ is consistent with PRMT1 being a host factor required for the virus.

SARS-CoV-2 viral protein extracts immunoblotted with ASYM26 revealed the arginine methylation of the N protein. It is known that the N protein is highly immunogenic and anti-N antibodies are amongst the first to appear in infected individuals ^71^. It is likely that SARS-CoV-2 N protein is fully methylated in infected cells, thus epitopes from the methylated RGG/RG peptides of N protein likely contribute to its elevated immunogenicity. Notably, R95- and R177-derived SARS-CoV-2 N peptides are predicted to be B cell epitopes ^72, 73^. Moreover, R95- and R177-peptides from the N protein were used to design SARS-CoV-2 vaccines in India ^72^, and we propose that the incorporation of asymmetrical dimethylarginines would increase their immunogenicity. The absence of methylation of the S protein observed using anti-ASYM26 is likely due to the lack of RGG/RG motifs. However, the S protein harbors an RRXR sequence within its furin cleavage site, a potential methylation sequence for PRMT7 ^74^. Thus, mass spectrometry analysis and various anti-methylarginine antibodies targeting MMA and SDMA will be needed to further define the sites of arginine methylation within the SARS-CoV-2 viral proteins.

Coronaviruses, like other viruses, have devised strategies to destabilize and inhibit SG formation to ensure optimal viral replication. The infectious bronchitis coronavirus (IBV) uses endoribonuclease Nsp15 for SG interference ^75^. For MERS-CoV, it is the 4a accessory protein that inhibits SG formation ^76, 77^. The N protein from SARS-CoV and SARS-CoV-2 were shown to localize to SGs ^14, 22, 26–28^. Thus, targeting the N protein function in G3BP1 SG regulation represents a new and valid strategy to fight COVID-19. It is known that the SR linker region when phosphorylated renders the N protein condensate more liquid-like and inhibition of N protein phosphorylation favors its translocation to SGs ^14, 18, 25, 29, 30, 78, 79^. We now show that arginine methylation of N protein is a post-translational modification that tunes N protein regulation of G3BP1 SGs. Information about how SGs are formed and regulated is emerging and represents a combination of multivalent interactions of protein-protein, protein-RNA and RNA-RNA interactions ^80^. How does arginine methylation of N protein regulate SG formation? Both R95K and R177K were defective in RNA binding to the SARS-CoV-2 5’-UTR RNA, and yet only R95K within the NTD was required for SG regulation. Our observation is in agreement with a recent report that residues 1-175 of N protein are sufficient to disrupt SGs ^26^. Consistent with R95 and R177 being part of the U-shaped β-platform of the RNA binding domain ^4, 11^, we show that both residues are needed to bind the 5’-UTR where the putative viral packaging signal of the genomic RNA resides. It is possible that N protein R177 is not needed to associate non-specifically with host mRNAs and hence its substitution to lysine does not influence SGs. Thus, we propose that RGG/RG motif methylation of the N protein affects SGs by modulating interaction with methylreaders and RNA.

The N interactome was not significantly altered in the presence of MS023, but there was an increase in PRMT1 association. We noted that increasing the concentration of a methylarginine reader TDRD3 blocked N protein from suppressing SG formation. Therefore, we propose that MS023 besides inhibiting the activity of PRMT1, also inactivates N protein by increasing its interaction with PRMT1, therefore allowing PRMT1 to become an enzyme-inactive RGG/RG motif reader. A role for arginine methylation of the RGG/RG motif in RBP phase separation is known ^32, 36^. For example, the RGG/RG motif protein FUS, dysregulated in ALS, undergoes liquid-liquid phase separation in the absence of PRMT1 ^81, 82^. Thus, arginine methylation is a key regulator of RNP condensation. As type I PRMT GSK3368715 inhibitor is in phase I clinical trials for diffuse large B-cell lymphoma ^41^, our data suggest that these inhibitors may be an effective strategy to interfere with N protein condensation and influence late stages of the SARS-CoV-2 life cycle.

Arginine methylation influences nearby phosphorylation sites often being antagonistic ^36^. For example, arginine methylation of the FOXO1 transcription factor at R248 and R250 by PRMT1 prevents AKT phosphorylation at S253, blocking nuclear exclusion of FOXO1 ^83^. Arginine methylation of cytoplasmic tail of the epidermal growth factor receptor at R1175 by PRMT5 enhances its trans-autophosphorylation at Y1173 ^84^. N protein RNA binding activity is known to be regulated by phosphorylation. *In vitro* studies showed that hypophosphorylation of the N protein facilitated interaction with RNA ^22^. We show that methylation of N protein R95 and R177 is needed for RNA binding. As S176, S180, S183, and S184 are reported to be phosphorylated by SRPK1 ^29^ and GSK3-Cdk1 ^18, 30, 78^, it is likely that there will be an interplay between phosphorylation and methylation especially near R177 for binding to the 5’-UTR of the SARS-CoV-2 RNA. It is likely that optimal binding of N protein to the 5’-UTR of the SARS-CoV-2 RNA will require a balance of methylarginines and phosphoserines. This interplay may also influence interactions with methylreaders including TDRD3 and phosphoreaders such as 14-3-3 proteins, the latter shown to bind the N protein ^85^.

In sum, our findings suggest that arginine methylation is required for N protein function, and PRMT1 is an essential regulator implicated in SARS-CoV2 life cycle. As PRMT inhibitors are in clinical trials ^36^, they may have applications for COVID-19, in addition to them being promising cancer drug candidates.

## Materials and Methods

### Reagents and antibodies

Immunoblotting was performed using the following antibodies: mouse anti-Flag monoclonal antibody (F1804, Sigma Aldrich, 1:2000); rabbit anti-SARS-CoV-2 N antibody (9103, Prosci, 1:2000); rabbit anti-SARS-CoV-2 S antibody (PA5-81795, Invitrogen, 1:1000); rabbit anti-TDRD3 antibody (Bethyl Laboratory, 1:1000); mouse anti-SMN antibody (610646, BD Biosciences, 1:2000); rabbit anti-G3BP1 antibody (1F1, Rhône-Poulenc Rorer, 1:1000, a kind gift from Dr. Imed Gallouzi at McGill University) ^86^; rabbit anti-ASYM26 (13-0011, Epicypher, 1:1000). Immunofluorescence were performed with the following antibodies: mouse anti-Flag monoclonal antibody (F1804, Sigma Aldrich, 1:500); rabbit anti-G3BP1 antibody (1F1, 1:500). Alexa Fluor-conjugated goat anti-rabbit, goat anti-mouse secondary antibodies were from Invitrogen. Protease inhibitor cocktail and protein phosphatase inhibitor cocktail were from Roche. MS023 (SML1555), sodium arsenite (S7400), Protein A-Sepharose (P3391) and PRMT5:MEP50 active complex (SRP0145) were purchased from Sigma Aldrich. Protein co-immunoprecipitation was performed using ANTI-FLAG® M2 Affinity Gel (A2220, Sigma Aldrich).

### Cell culture and transfection

HEK293 and VeroE6 cells were maintained in Dulbecco’s modified Eagle’s medium (DMEM) supplemented with 10% fetal bovine serum (FBS) and grown at 37°C with 5% CO_2_. Huh-7 cells were maintained in Dulbecco’s modified Eagle’s medium (DMEM) supplemented with 10% fetal bovine serum (FBS) and Non-Essential Amino Acid (NEAA, Gibco) and grown at 37°C with 5% CO2. Cells were transfected with 20 nM siRNA oligonucleotides using Lipofectamine RNAiMAX (Invitrogen) according to the manufacturer’s instructions. HEK293 and Huh-7 cells were transfected with plasmid DNAs by standard calcium phosphate precipitation and Lipofectamine 3000 (Invitrogen), respectively.

### Plasmids and siRNAs

The N-terminal Flag-tagged SARS-CoV-2 N plasmid was constructed by inserting a Flag-coding sequence into the pcDNA3.1 (+) vector at the *Hind* III and *Bam* HI sites to generate pcDNA3.1-Flag and then the PCR-amplified cDNA of SARS-CoV-2 N coding region at *Bam* HI and *Xho* I sites of pcDNA3.1-Flag vector. The PCR template DNA is a plasmid with insertion of synthesized DNA expressing SARS-CoV-2 N protein provided by Dr. Shan Cen based on the SARS-CoV-2 Wuhan-Hu-1 isolate (GenBank: MN_908947). The plasmids for expressing GST fusion proteins of SARS-CoV-2 N truncated fragments were constructed by inserting PCR-amplified SARS-CoV-2 N cDNA fragments in pGEX-6P1 vector at *Bam* HI and *Sal* I sites. The GST-RGG construct was generated by inserting a DNA fragment expressing the mouse RBMX C-terminal RGG/RG motif in pGEX-6P1 vector at *Bam* HI and *Sal* I sites. The mutants with replacement of the arginine residues with lysine at the RGG/RG motif were constructed by a two-step PCR strategy. p5’-UTR:CoV-2 was constructed using synthesized DNA fragment with the 1-400 bp of 5’-UTR of SARS-CoV-2 gRNA (NC_045512) and the DNA fragment was insert into pcDNA3.1 vector at *Bam* HΙ and *Xba* Ι sites. The myc-tagged SMN and TDRD3 plasmids were generated in previous studies ^87, 88^. All siRNAs were purchased from Dharmacon. siRNA sequences are as follows: siPRMT1, 5’-CGT CAA AGC CAA CAA GTT AUU-3’. The siRNA 5′-CGU ACG CGG AAU ACU UCG AdTdT-3′, targeting the firefly luciferase (GL2) was used as control. 20 nM siRNA was used for transfection.

### Protein purification and *in vitro* methylation assay

Expression of GST fusion proteins in bacteria was induced with 1mM isopropyl-β-D-thiogalactopyranoside (IPTG) at room temperature for 16 h. All steps of the purification after growth of bacteria were performed at 4°C. Cells were lysed by sonication in PBS buffer containing a mixture of protease inhibitors. The lysate was clarified by centrifugation and the supernatant was incubated with glutathione-Sepharose 4B beads for 2 h. The resin was washed four times with PBS buffer and then twice with 50 mM Tris-HCl pH7.4 buffer. Protein was eluted with 10 mM reduced glutathione in 50 mM Tris-HCl pH 7.4 buffer. Approximately 10 μg of each GST fusion protein was incubated with 1 μl of (methyl-^3^H) S-adenosyl-L-methionine solution (15Ci/mmol stock solution, 0.55 μM final concentration, Perkin-Elmer) and 1-2 μg of PRMTs in methylation buffer (50mM Tris-HCl pH 7.4, 1 mM DTT) for 1 to 4 h at 25°C or 37°C. Samples were separated by SDS-PAGE and stained with Coomassie Blue. After de-staining, the gel was then incubated for 1 h in EN^3^HANCE (Perkin Elmer) according to manufacturer’s instructions and the reaction was visualized by fluorography.

### Cell lysis, immunoprecipitation, immunoblotting and LC-MS/MS analysis

For co-immunoprecipitation experiments, cells were lysed in lysis buffer (50 mM HEPES, pH 7.4, 150 mM NaCl, 1% Triton X-100 and a cocktail of protease inhibitors and phosphatase inhibitors). Cell lysates were cleared with high-speed centrifugation to remove cell debris, then the supernatant was incubated with anti-Flag M2 beads for 1.5 h at 4°C. Samples were washed with 1 ml of lysis buffer for 3 times and eluted with 2× SDS loading buffer for western blot analysis. For LC-MS/MS, the beads were further washed with PBS buffer twice. The beads together with the bound proteins were subjected to LC-MS/MS performed at the mass spectrometry facility at IRIC, Université de Montréal. The data were analyzed using Scaffold Proteome Software.

### MS023 toxicity assay

Prior to performing viral infections, the MS023 inhibitor was examined for toxicity to the VeroE6 cells. VeroE6 cells (2,500 cells per well) were seeded in 96-well plate and cultured at a condition similar to that in the viral infection analysis. The type I PRMT inhibitor MS023 was dissolved in DMSO and diluted in complete DMEM containing 10% FBS. The medium was added to cells with a final concentration of 2 to 30 μM MS023 and 0.2% DMSO as indicated. Twenty-four hours later, the cell culture medium was replaced with 2% FBS/DMEM medium containing the same concentration of MS023 and DMSO in the corresponding wells. After 48 h of further incubation, the cell viability was analyzed using the MTT assay kit (Abcam, ab211091) according to manufacturer’s instructions. Briefly, the media was carefully removed, and both 50 μl of serum-free medium and 50 μl of MTT reagent were added to each well. Following 3 h of incubation at 37°C, the MTT reagent was removed and 150 μl of MTT solvent was added to each well and the plate was incubated at room temperature on an orbital shaker for 30 min prior to reading absorbance at OD_590_ nm. The absorbance of MS023-treated wells was divided by the absorbance of the DMSO-treated wells to normalize cell survival.

### SARS-CoV-2 infection and purification of genomic SARS CoV-2 RNA (gRNA) and proteins

Within the certified BL3 containment facility of the McGill University Health Centre, VeroE6 cells were seeded in 24-well plates (10^5^/well in 0.5 ml) and incubated in complete DMEM containing 10% FBS in the presence of PRMT1 inhibitor MS023 or DMSO control for 24 h prior to infection. The cells were then infected with SARS-CoV-2 isolate RIM-1 (GenBank accession number: MW599736) at multiplicity of infection (MOI=0.1) at 37°C for 2h. The virus inoculum was removed, and the cells were washed once with 2% FBS/DMEM medium and then incubated for an additional 48 h in 1 ml of 2% FBS/DMEM containing the PRMT inhibitor at indicated concentrations or same amount of DMSO as control at 37°C. After the infection was complete, 250 μl cell supernatant was lysed in 0.75 ml TRIzol LS (Invitrogen) and transported out of the BL3 facility. The viral RNA was then extracted from the TRIzol using chloroform extraction following manufacturer’s instructions. One step RT-qPCR was performed using TaqMan™ Fast Virus 1-Step Master Mix following the manufacturer’s instructions. Viral gRNA was detected using primers (Fw: 5’-ATG AGC TTA GTC CTG TTG-3’, Rv: 5’-CTC CCT TTG TTG TGT TGT-3’), and probe (5’Hex-AGA TGT CTT GTG CTG CCG GTA-BHQ-1-3’), specifically targeting *RdRp* gene as described ^89^. In addition, viral proteins were extracted from the organic phase of TRIzol solution according to the manufacturer’s instruction. Briefly, after removing the aqueous phase, 0.3 ml 100% ethanol was added to the organic phase. Genomic RNA (gRNA) from infected cells was pelleted by centrifugation at 2000g for 5 min. 0.75 ml supernatant was moved to a new tube and mixed with 1.5 ml isopropanol. Proteins were collected by centrifugation at 12000 rpm for 15 min, following by two times of washing with 0.3 M guanidine hydrochloride and 95% ethanal. Liquid was removed and the pellet was air-dried. The dried proteins were dissolved in 1XSDS loading buffer and proceed with western blot analysis.

### PAR-CLIP (photoactivatable ribonucleoside-enhanced crosslinking and immunoprecipitation) and RNA immunoprecipitation

PAR-CLIP (photoactivatable ribonucleoside-enhanced crosslinking and immunoprecipitation) was performed as previously described with minor modification ^90^. p5’-UTR:CoV-2 and pFlag-N were co-transfected to HEK293 cells with a 1:1 ratio. 24 h post-transfection, the cells were treated with 100 μM 4-thiouridine for 16 h and cross-linked with 0.15 J/cm^2^ 365 nm UV. Cells were then washed twice with ice cold PBS and resuspended in lysis buffer (150mM KCl, 25mM Tris, pH 7.4, 5mM EDTA, 0.5mM DTT, 0.5% NP40, and 100U/mL RNase inhibitor). After incubation for 20 min with rotation at 400 rpm and cell debris were cleared by centrifugation. Cell lysates were incubated with 1 U/ml RNase I at 37 °C for 3 min. For each immunoprecipitation, 40 U RNase inhibitor and 2 μg of antibody was added, and the samples were incubated for 2 h at 4°C with rotation. Protein A Sepharose beads (Sigma) were then added and the samples were incubated at 4°C for another hour with rotation. The beads were pelleted by centrifugation, resuspended, and washed in high salt wash buffer for 3 times and lysis buffer once. After removing the final wash buffer, DNA fragments were digested with 2U TURBO™ DNase (Thermo fisher, AM2238) at the 37°C for 4 min. RNA was eluted in Proteinase K buffer (50 mM Tris, pH 7.5, 75 mM NaCl, 6.5 mM EDTA, and 1% SDS) supplemented with proteinase K and incubated at 50°C for 30 min. RNA was recovered by using 5 volumes of TRIzol™ LS Reagent (Thermo Fisher). Equal volume of RNA from each sample was used for the reverse transcription. qPCR was performed using primers targeting gRNA 5’-UTR. Primer sequences used in the experiment are as follows: position 1: Fw: 5’-TCG TTG ACA GGA CAC GAG TA-3’, Rv: 5’-CCC GGA CGA AAC CTA GAT GT-3’; position 2: Fw: 5’-CCT TGT CCC TGG TTT CAA CG-3’, Rv: 5’-CAC GTC GCG AAC CTG TAA AA-3’. RNA immunoprecipitation (RIP) was performed as previously described with minor modifications ^91^. Briefly, cells were cross-linked with a final concentration of 1% formaldehyde, washed twice with ice cold PBS and resuspended in RIP buffer (150mM KCl, 25mM Tris, pH 7.4, 5mM EDTA, 0.5mM DTT, 0.5% NP40, and 100U/mL RNase inhibitor). Chromatin was sheared by sonication, and DNA fragments were digested with TURBO™ DNase (Thermo fisher, AM2238) at the 37°C for 15 min. Cell lysate were proceeded with immunoprecipitation with antibody and wash with RIP buffer. The RNA was recovered from the precipitate as described above. qPCR was performed using primers: Fw: 5’-TCG TCT ATC TTC TGC AGG CT-3’, Rv: 5’-ACG TCG CGA ACC TGT AAA AC-3’.

### Arsenite treatment and immunofluorescence

Cells growing on glass coverslips were treated with 0.5 mM arsenite for 1h and fixed for 10 min with 4% paraformaldehyde (PFA). After three washes with PBS the cells were permeabilized for 5 min with 0.25% Triton X-100 in PBS. Coverslips were incubated with blocking buffer containing 5% FBS for 1h, and then incubated with primary antibodies diluted in PBS containing 5% FBS for 2h. After three washes, the coverslips were incubated with corresponding fluorescent secondary antibodies for another hour in PBS buffer containing 5% FBS. After rinsing, the coverslips were mounted with IMMU-MOUNT (Thermo Scientific) mounting medium containing 1µg/ml of 4′,6-diamidino-2-phenylindole (DAPI). Images were taken using a Zeiss LSM800 confocal microscope.

### Statistical analysis

All data are expressed as mean ± S.E.M. and compared between groups using the Welch’s t test. *p* value <0.05 was considered statistically significant. *, *p* < 0.05; **, *p* < 0.01; ***, *p* < 0.001.

## Acknowledgements

We thank Dr. Marcel Behr, Fiona McIntosh and Andreanne Lupien from the Research Institute of McGill University Health Centre for access to the biosafety level 3 lab, providing SARS-CoV-2 strain RIM-1, and expert advice. We thank Dr. Imed Gallouzi for anti-G3BP1 antibodies and Dr. Shan Cen for providing SARS-CoV2 N plasmid DNA. This work was funded by a Canadian Institute of Health Research FDN-154303 award to S. R. T.C. holds a Fonds de recherche du Québec en Santé studentship award.

## Author Contributions

T.C., Z.Y, C.L and S.R. designed the research; T.C., Z.Y and Z.W. performed the experiments; T.C., Z.Y, and S.R. analyzed the data; T.C., Z.Y, C.L and S.R. wrote the paper.

